# Germline RNA helicases couple RNA binding to P granule assembly at nuclear periphery

**DOI:** 10.1101/760256

**Authors:** Wenjun Chen, Yabing Hu, Charles Lang, Jordan S Brown, Xiaoyan Song, Ed Munro, Karen Bennett, Donglei Zhang, Heng-Chi Lee

## Abstract

P granules are phase-separated liquid droplets that play important roles in the maintenance of the germ cell fate in *C. elegans*. The localization and formation of P granules are highly dynamic, but mechanisms that regulate such processes remain poorly understood. Here we show that germline RNA helicases (GLHs) control the formation and disassembly of germ granules through their binding and release of RNAs, respectively. In addition, the FGG repeats in the GLHs promote the formation of germ granules at the perinucleus. Proteomic analyses of a mutation that traps RNA-bound GLH-1 complex revealed transient interactions of GLH-1 with several Argonautes and RNA binding proteins. Finally, we found that defects in perinuclear P granule formation correlate with the fertility defects observed in various GLH mutants. Together, our results highlight the versatile roles of RNA helicases in controlling the formation of liquid droplets in space and time.

## INTRODUCTION

Germ granules, also known as P granules in *C. elegans*, are phase separated liquid droplets that are frequently found at the perinuclear region in the germ cells of all animals (Brangwynne et al., 2009; Eddy, 1975). In *C. elegans*, P granules have been shown to function in maintaining germ cell fate (Updike et al., 2014). The formation and localization of P granules are dynamic but tightly regulated during germ line development. P granules are first assembled in the cytoplasm of one-cell embryos, where they are sorted asymmetrically to cells of the germ line lineage. Starting at the four-cell stage, P granules start to anchor to the nuclear periphery of the germ cell precursor, where they stay throughout most of germ cell development (Updike and Strome, 2010). RNA binding proteins such as GLH-1, PGL-1 and MEG-3/MEG-4 have been reported to play essential roles in the formation of P granules (Hanazawa et al., 2011; Updike and Strome, 2009; Wang et al., 2014). While *pgl-1* and *glh-1* single mutants exhibit defects in the formation of both cytoplasmic and perinuclear P granules, the *meg-3/meg-4* double mutant animals only exhibit defects in the formation of cytoplasmic P granules of embryos. Recent studies have shown that the gradients of RNA bound PGL-1 and MEG-3/MEG-4 are responsible for the formation of germ granules at the posterior of one-cell embryos (Saha et al., 2016; Smith et al., 2016). However, much less is known about how GLH-1 regulates germ granule formation, or how the formation of perinuclear P granules is controlled.

GLH-1 is one of the VASA helicase homologues in *C. elegans* and was the first constitutive component of P granules identified (Roussell and Bennett, 1993)(Gruidl et al., 1996). Subsequently three additional GLH family members (GLH-2, GLH-3 and GLH-4) were reported (Kuznicki et al., 2000a). VASA mutants exhibit fertility defects in all animals tested so far (Gustafson and Wessel, 2010). Similarly, depletion or deletion of *glh* genes produce reduced broods and sterile progeny (Kuznicki et al., 2000b; Spike et al., 2008), and the loss of both GLH-1 and GLH-4 results in complete sterility. All the GLHs belongs to the DEAD box family of RNA helicases and have the requisite RNA helicase motifs. GLH-1,2, and 4 also possess distinct N-terminal Phenylalanine-Glycine-Glycine (FGG) repeats. While the function of these FGG repeats in GLH-1 is not known, it was proposed that FGG repeats may facilitate the interaction of GLH-1 with the FG-hydrogel formed at nuclear pores (Frey et al., 2006; Updike et al., 2011a). In support of this, knockdown of many nuclear pore factors leads to loss of perinuclear germ granules (Updike and Strome, 2009; Voronina et al., 2010). In addition, a GFP fusion protein that contains multimeric FGG repeats fused to the N terminus of GFP leads to formation of some GFP granules at the nuclear periphery (Updike et al., 2011a). However, the direct evidence that FGGs in GLH-1 tethers P granules to the nuclear periphery remains missing. In addition, it was unclear whether the ATPase/RNA helicase activity, and/or the RNA binding activity of GLH-1 is required for its role in P granule formation.

Here we show that GLH-1 promotes the assembly and disassembly of germ granules through its binding and release of RNAs, and these processes are coupled to distinct steps of the GLH-1 ATP hydrolysis cycle. In addition, the FGG repeats in the GLHs are required for the robust formation of germ granules into perinuclear foci. Proteomic analyses suggest that RNA-bound GLH-1 complexes concentrate Argonautes and other RNA binding proteins at the nuclear periphery. Our analyses support a model that GLHs control the dynamics and nuclear anchoring of P granules to promote germ cell development.

## RESULTS

### GLH helicases promote the assembly of P granules through their RNA binding

To investigate the role of the GLH-1 RNA helicase in P granule formation and the localization of piRNA pathway factors, we used GFP-tagged PRG-1, a PIWI Argonaute enriched in P granules, as a P granule marker and examined its localization in *glh-1* mutants (Batista et al., 2008). Consistent with previous reports, a *glh-1* null mutation reduced both size and number of P granules (Figure 1A). As it was reported that GLH-1 and GLH-4 act redundantly to promote fertility in *C. elegans* (Kuznicki et al., 2000b; Spike et al., 2008), we suspected that GLH-1 and GLH-4 may play redundant roles in P granule formation and therefore examined the PRG-1 granules in the *glh-1glh-4* double mutant. While P granules look normal in a *glh-4* mutant, we observed little, if any, PRG-1 granules in *glh-1glh-4* double mutants (Figure 1A). Similar results were observed using GFP::PGL-1 as a marker of granules (Figure S1A). Together, these observations suggest GLH-1 and GLH-4 jointly promote P granule formation, but GLH-1 plays a more major role in this process.

**Figure 1.**
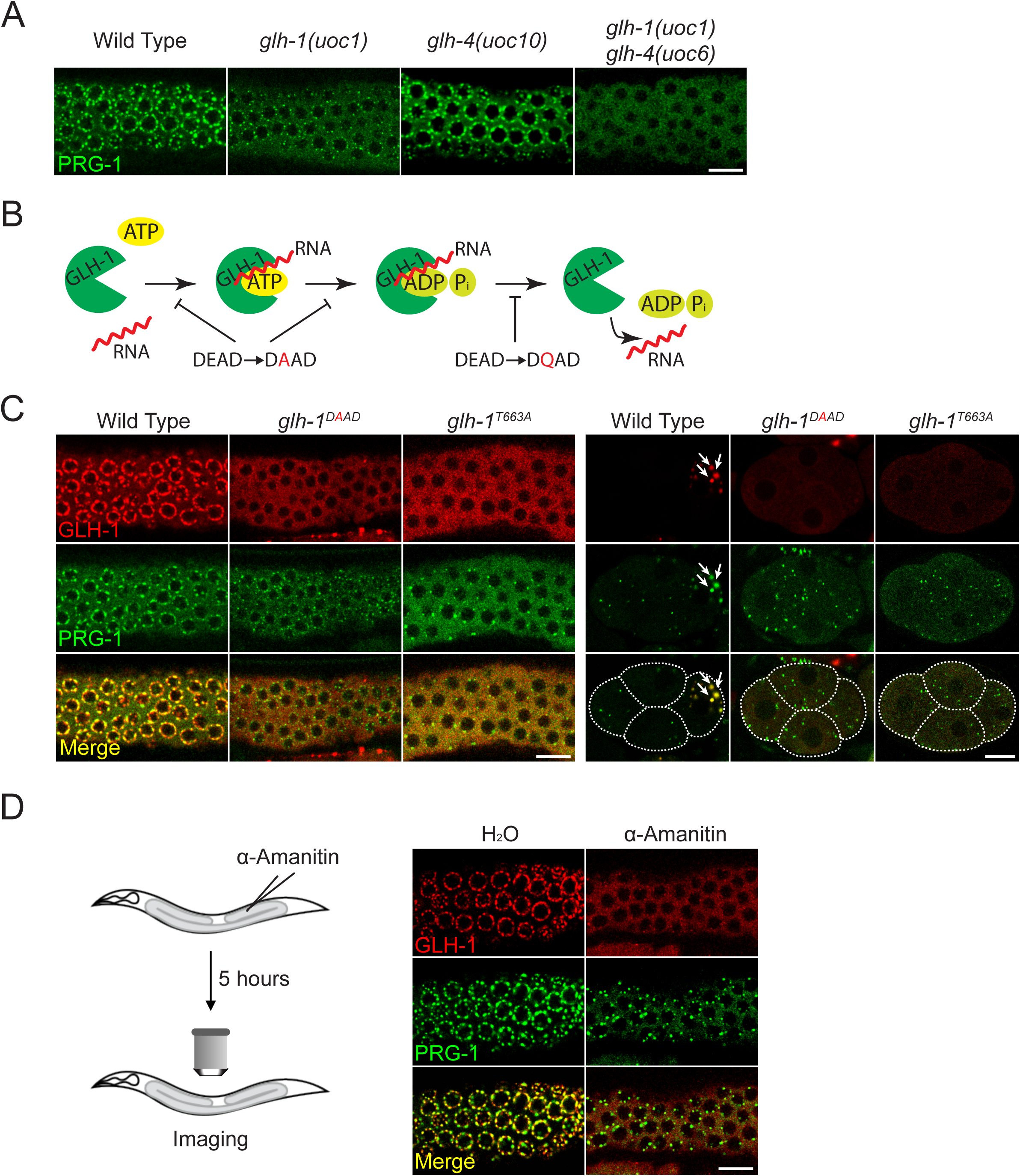
Germline helicases promote the formation of P granules through its RNA binding. (A) GLH-1 and GLH-4 act redundantly to promote the formation of P granules. Fluorescent micrographs showing the localization of P granule component GFP::PRG-1 in the indicated strains. Note that *uoc1, uoc6* and *uoc10* are gene-edited null alleles of the indicated genes. (B) A model showing how GLH-1 couples its ATP hydrolysis cycle to its RNA binding/release and the proposed effects of indicated mutations (C) The localization of mCherry::GLH-1 and GFP::PRG-1 in the indicated strains in the adult germline (left) or in the 4-cell embryo (right). Arrow indicates the cytoplasmic P granules in the 4-cell embryo. (D) P granules are dispersed into the cytoplasm 5 hours after the injection of α-Amanitin, an RNA polymerase II inhibitor.

In addition to the well-characterized role of DEAD box helicases in unwinding short double strand RNAs, the GLH homologue VASA has been proposed to act as an ATP-dependent RNA clamp, where ATP binding is coupled to RNA binding (Xiol et al., 2014). As RNA binding by PGL-1 and MEX-3/4 plays a critical role in P granule formation (Saha et al., 2016; Smith et al., 2016), we wondered whether RNA binding of GLH helicases contributes to their role in P granule assembly. We created gene-edited worms that express GLH-1 with specific mutations that affect distinct steps of the ATP hydrolysis cycle of DEAD box helicases (Figure 1B). We first examined the effects of a DAAD mutation, which changes glutamic acid in DEAD box to alanine. This mutation is expected to abolish the ability of the DEAD box to bind and hydrolyze ATP, and in many cases also reduces RNA binding activity (Putnam and Jankowsky, 2013). In transgenic worms expressing GLH-1 with a DAAD mutation, we observed a reduction in the size and number of P granules (Figure 1C left). Similarly, very few P granules are found in early embryos of the GLH-1 DAAD mutant, where *de novo* assembly of P granules occurs (Figure 1C, right). This observation suggests that the ATP binding and likely the RNA binding of GLH-1 promotes the assembly of P granules. To further examine the role of RNA binding of GLH-1 in P granule formation, we examined the effects of a mutation in the conserved threonine at motif V of GLH-1 helicase domain (T663A), which is specifically involved in RNA binding of VASA helicases by directly contacting the RNA backbone (Sengoku et al., 2006). We observed similar granule defects in the *glh-1* T663A mutant as in the *glh-1* DAAD null mutant, suggesting that the RNA binding of GLH-1 promotes P granule assembly (Figure 1C). In addition, we examined the effect of RNase on P granules *ex vivo*. We reasoned that if RNAs play an important role in P granule assembly, RNase should destabilize P granules. Indeed, after extruding the adult germline from the body cavity, we observed that all P granules dissipated in RNase-containing buffer within two minutes, while around 50 percent and 25 percent of P granules remain visible after two and five minutes in control buffer, respectively (Figure S1B). Consistent with a previous report (Sheth et al., 2010), we also observed that injection of α-amanitin, an RNA Polymerase II inhibitor, into the adult germline leads to the disassembly of P granules (Figure 1D). This observation suggests that the maintenance of P granules requires newly transcribed RNAs. Together, our results suggest that the RNA binding activity of the GLH-1 helicase is required for the assembly of P granules.

### GLH helicases require RNA release to promote the disassembly of P granules

Interestingly, in worms expressing a GLH-1 DQAD mutation, a mutation which is shown to interfere with the release of ATP hydrolyzed products and the release of RNAs (Xiol et al., 2014)(Figures 1B), P granules are drastically enlarged (Figure 2A and Figure S2A). Notably, the majority of these enlarged granules found in the DQAD mutants have lost their perinuclear localization in the germ cells of the adult germline. We reason that the enlarged, untethered granules are caused by defects in P granule disassembly. If so, we expected the exchange of germ granule factors between cytoplasmic pools and these germ granules would be reduced. Indeed, using a fluorescent recovery after photobleaching assay (FRAP), we observed that while the P granule components GLH-1 and PRG-1 exhibit significant recovery of fluorescent signals within 10 seconds after photobleaching in the wild type animals, their recovery is significantly reduced in the DQAD mutant (Figures 2B-D and Figure S2B). PRG-1 exhibits a slightly faster and greater recovery than GLH-1 in the GLH-1 DQAD mutant, likely due to the function of other GLH helicases in regulating PRG-1 dynamics. Since the asymmetric sorting of P granules relies on the differential rate of assembly/disassembly of P granules between anterior vs. posterior (Wang et al., 2014), we expected these enlarged, less dynamic granules would exhibit defects in P granule sorting. Indeed, unlike in wild type animals, in GLH-1 DQAD mutants, P granules are not sorted to the germline lineage, but rather are randomly distributed to cells of both somatic and germline lineages in early embryos (Figure 2A, right and movie 1&2). Furthermore, we reasoned that if these enlarged P granules are caused by defects in RNA release from GLH-1, disruption of GLH-1 RNA binding should interfere with aggregation. Indeed, in *glh-1* mutants carrying both the T663A and DQAD mutations, we no longer observed enlarged P granules, but rather saw reduced P granules as seen in *glh-1* T663A mutants alone (Figure 2A). This observation suggests that RNA binding is required for the GLH-1 DQAD mutant to form enlarged P granules. Together, our analyses provide a simple model of how the dynamics of P granules are controlled: GLHs promote the assembly and disassembly of P granules by binding and releasing RNAs, respectively.

**Figure 2.**
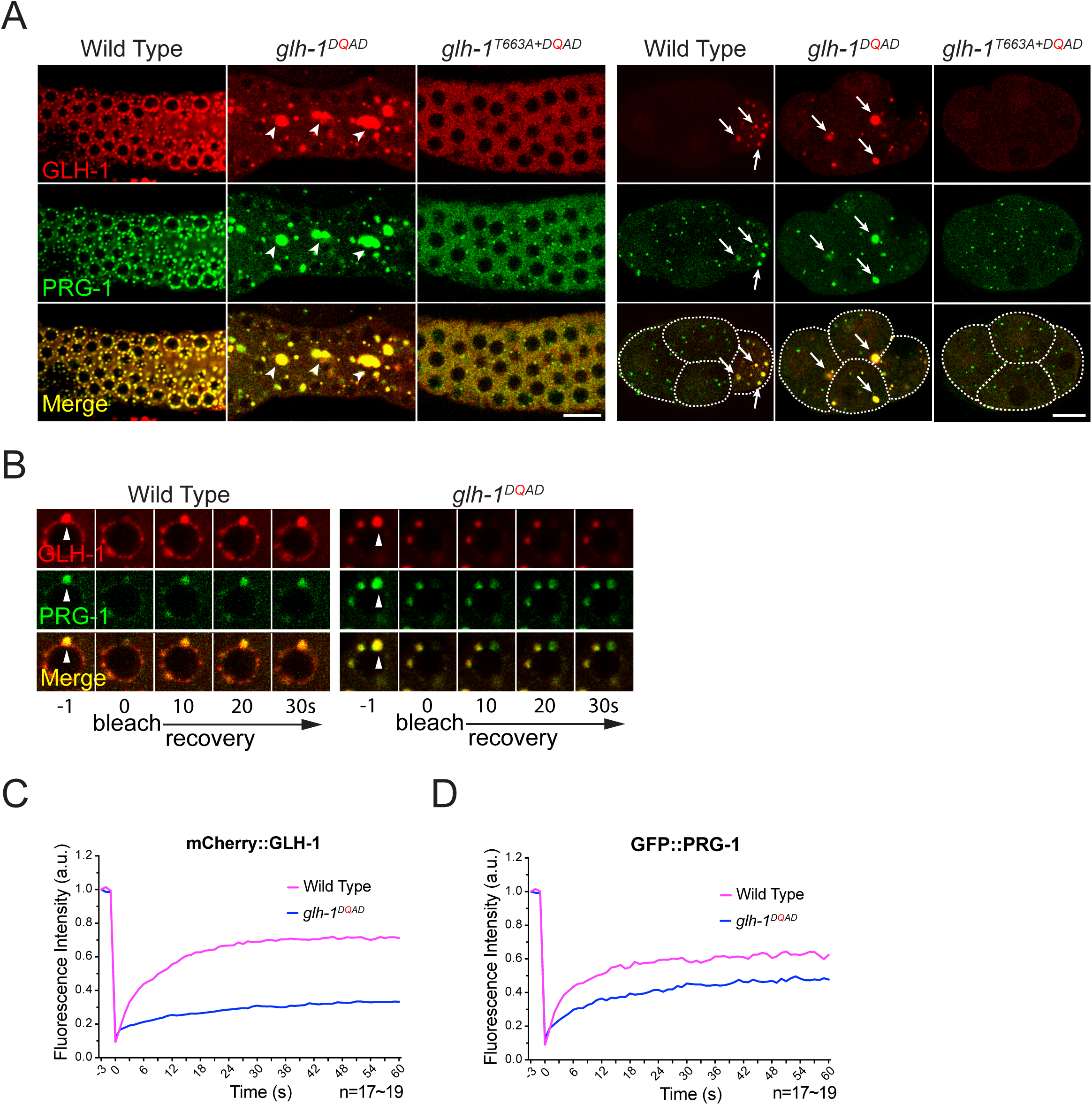
Germline helicases couple RNA release to promote the disassembly of P granules. (A) The localization of mCherry::GLH-1 and GFP::PRG-1 in the indicated strains in the adult germline (left) or in the 4-cell embryo (right). The arrowheads indicate many of the enlarged aggregates of P granules in the adult germline do not associate with germline nuclei. The arrows indicate in embryos, P granules are properly sorted to the cell of germline lineage (P cell) in wild type but are improperly sorted into cells of the somatic lineage in *glh-1* DQAD mutant. (B) Fluorescent Recovery After Photobleaching (FRAP) analyses indicate reduced dynamics of P granule components, including GLH-1 and PRG-1, in the adult germlines of *glh-1* DQAD mutant worms. (C-D) The quantification of FRAP analyses of (D) showing the average fluorescent signals of mCherry::GLH-1 (E) and GFP::PRG-1 (F) at the indicated times (seconds after photobleaching).

### RNA bound GLHs drive P granule assembly which enriches Argonaute proteins

We reasoned that since the GLH-1 DQAD mutation significantly reduces the disassembly of germ granules, it offered an opportunity to isolate the components of germ granules that otherwise may be too unstable to capture. Indeed, a previous study has found that the DQAD mutation in VASA leads to the stabilization of transient interactions between VASA and its interactors, including the PIWI Argonautes and factors required for piRNA amplification in insect culture cells (Xiol et al., 2014). We therefore performed proteomic analyses to examine the components of the GLH-1 and PIWI PRG-1 complexes in various GLH mutant backgrounds (Figure S3). In wild type GLH-1 background, we identified few, if any, GLH-1 and GLH-4 peptides in PRG-1 immunoprecipitated complexes (Figure 3A), implying the interactions between PRG-1 and GLH1/4 are transient. On the contrary, we identified significantly more GLH-1, GLH-2, and GLH-4 peptides in PRG-1 complex in the GLH-1 DQAD mutant background than in wild type animals (Figure 3A), suggesting the interaction between PRG-1 and these GLHs are stabilized. Similarly, in GLH-1 immunoprecipitated complexes, we identified significantly more peptides of germline-enriched Argonautes, including PRG-1, WAGO-1, WAGO-4, CSR-1 and the polyA binding protein PAB-1 in the GLH-1 DQAD mutant background than in wild type animals (Figure 3B). The interaction between GLH-1 and PRG-1 in the DQAD mutant background was further confirmed by Western blot analyses (Figure 3C). Since many GLH-1 interacting factors are known to function in gene silencing pathways (Flamand et al., 2016; Gu et al., 2009; Wan et al., 2018), our observations suggest that GLHs and P granules may facilitate small RNA-mediated gene silencing. Notably, the interactions between GLH-1 with various Argonautes can only be detected in GLH-1 complexes of DQAD mutants, while the interactions between GLH-1 with translation factors, including EFT-3 and EEF-2 are found in various GLH backgrounds, including in wild-type animals, DAAD and DQAD mutants. Since the release of RNA from GLH-1 is expected to be defective in the DQAD background, these results imply that the transient interactions between GLH-1 and Argonautes are likely mediated through RNAs. Together, these observations suggest a model that RNA bound GLH-1 drives the assembly of P granules, which are enriched in Argonautes and other RNA binding proteins.

**Figure 3.**
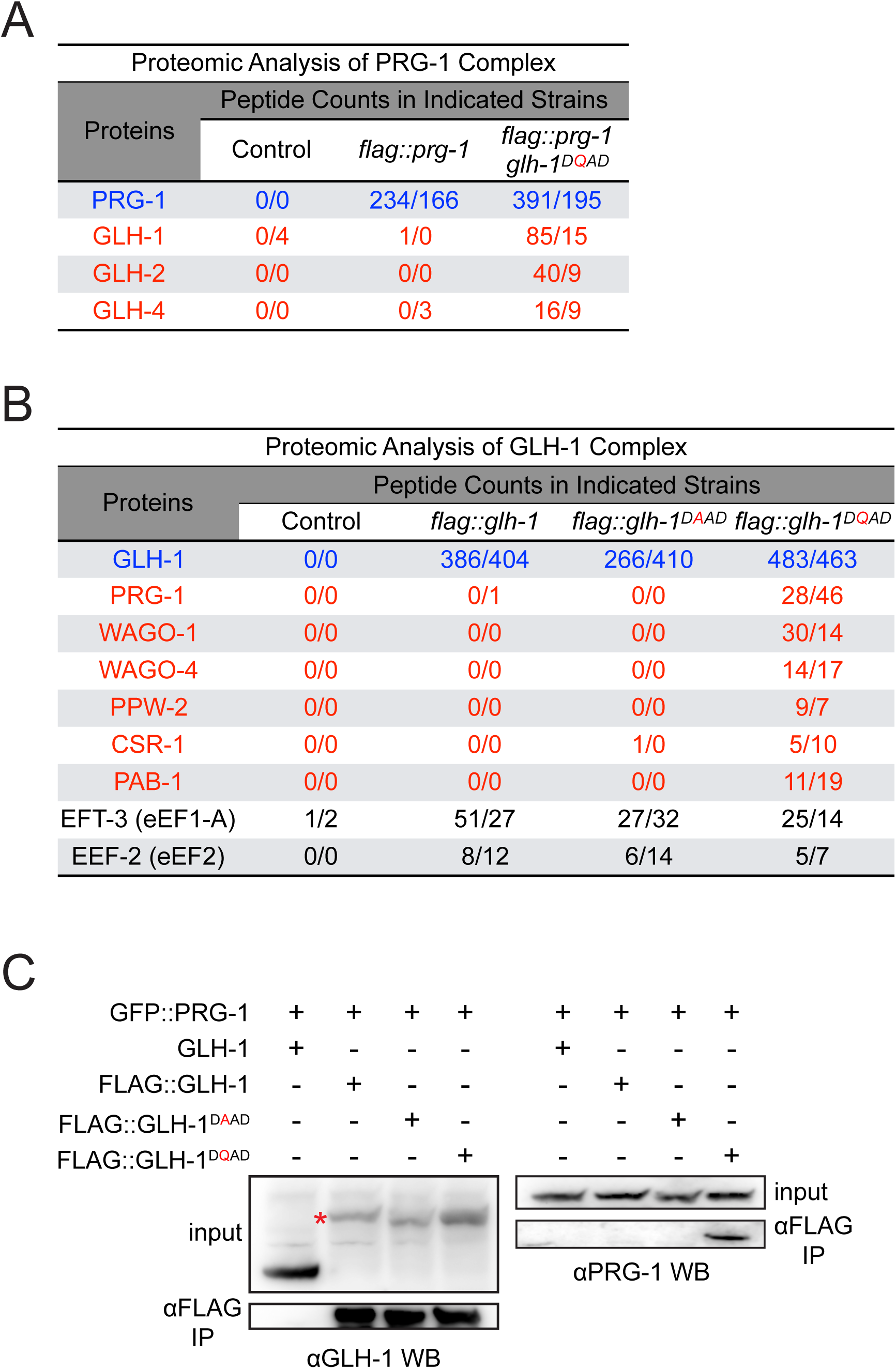
The RNA bound GLH-1 forms a complex that concentrates Argonautes and other RNA binding proteins. (A) Proteomic analyses of the PIWI PRG-1 complex in the indicated strains, wild type *glh-1* and *glh-1* DQAD mutant. The numbers of peptides of the indicated proteins identified in two independent experiments are shown. (B) Proteomic analyses of the GLH-1 complex in the indicated *glh-1* mutant backgrounds. The proteins whose GLH-1 interactions are stabilized in the *glh-1* DQAD mutant are highlighted in red. The numbers of peptides of the indicated proteins identified in two independent experiments are shown. (C) Western blot analyses showing the interaction between mCherry::GLH-1 and GFP::PRG-1 is stabilized in the *glh-1* DQAD strain. The red asterisk indicates the FLAG::GLH-1.

### The FGG repeats in GLH-1 promote perinuclear P granule localization

Our results suggest that P granule dynamics can be controlled by GLHs, but the mechanism by which P granules localize to the nuclear periphery remains unknown. Germ (P) granules are frequently found at the nuclear periphery in various animals (Eddy, 1975) and previous electron microscopy analyses suggest that germ granules localize to the cytoplasmic face in proximity to clusters of nuclear pores in *C. elegans* (Sheth et al., 2010). How germ granules anchor to the nuclear periphery remains largely unknown. As mentioned in the introduction, one intriguing model has been proposed that FGG repeats of the GLHs interact with the FG meshwork of nuclear pore proteins and thus help anchor germ granules to nuclear pore clusters (Updike et al., 2011b). To directly test the role of the GLH-1 FGG repeats in P granule formation, we examined the germ granules from worms expressing a GLH-1 mutant protein with its N terminus deleted, which removes all of its FGG repeats (*glh-1* FGGΔ). In early embryos, no obvious defects of cytoplasmic P granules are found in the *glh-1* FGGΔ mutant, nor in *glh-1* FGGΔ in the *glh-4* null background, suggesting the FGG repeats of GLH-1 are not required for the formation of cytoplasmic P granules in embryos (Figure 4A and Figure S4A). However, in the adult germline, perinuclear GLH-1 is significantly reduced in *glh-1* FGGΔ or in *glh-1* FGGΔ *glh-4* double mutants, while the perinuclear PRG-1 granules are partially reduced (Figure 4A and Figure S4A). The degree of perinuclear P granule defects are variable in *glh-1* FGGΔ mutant, but such defects were further exacerbated when worms were grown at a higher temperature, where very few perinuclear P granules are found in *glh-1* FGGΔ (Figure 4A and S4B). Overall we noticed the defects of perinuclear localization in *glh-1* FGGΔ is more dramatic for GLH-1 itself than PRG-1, suggesting additional GLHs or other unknown factors can promote the formation of some residual PRG-1 containing granules at perinuclear foci in *glh-1* FGGΔ mutants. Taken together, these observations indicate that the FGG repeats of GLH-1 contribute to the formation of perinuclear P granules, likely through anchoring GLH-1 itself to the FG hydrogel formed at nuclear pores.

**Figure 4.**
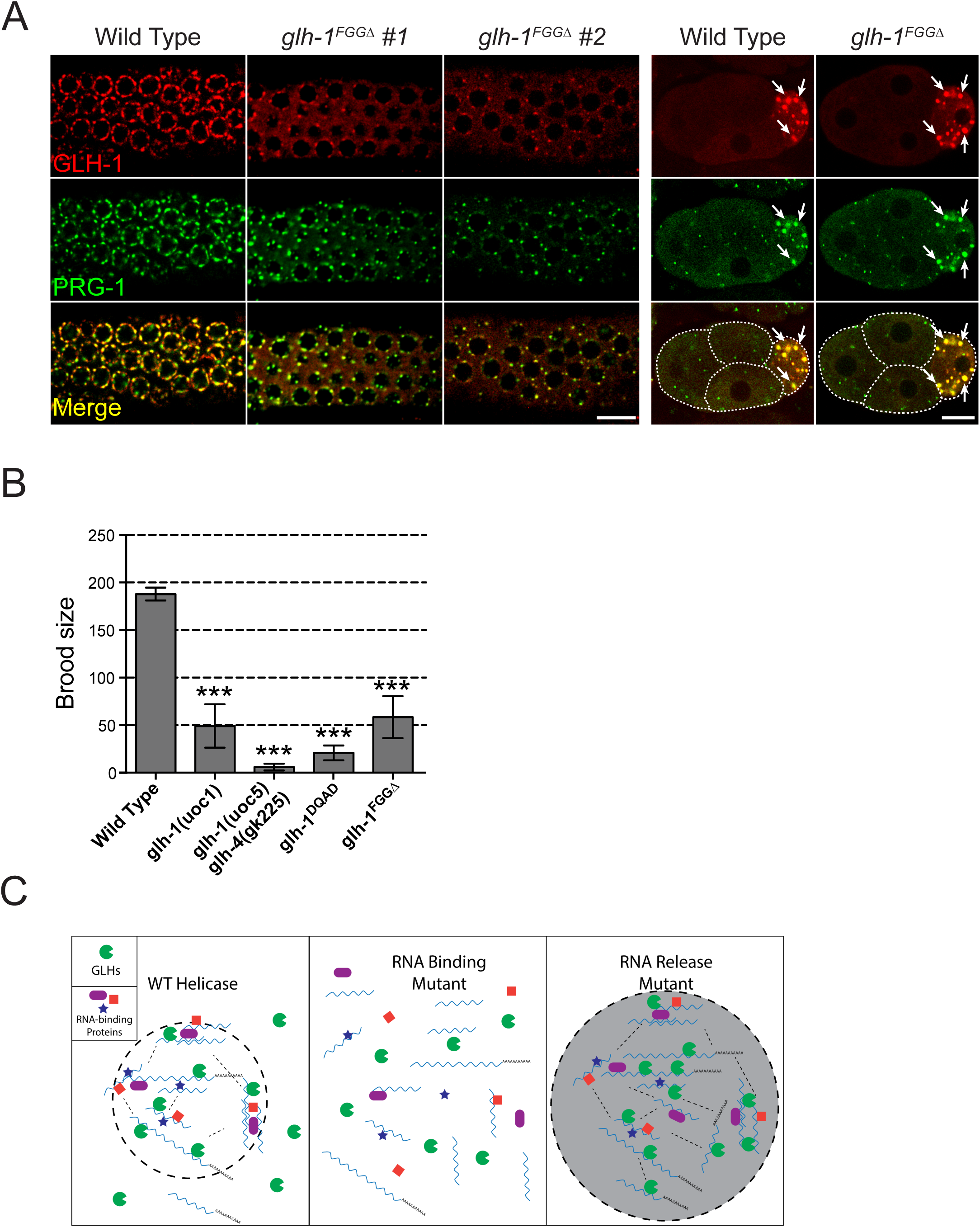
The N terminal FGG repeats in GLH-1 promote nuclear anchoring of P granules which is critical for fertility. (A) The formation of perinuclear P granules in the adult germline (left), but not of cytoplasmic P granules in embryos (right), is compromised in the *glh-1*^FGGΔ^ mutant. Images showing the localization of mCherry::GLH-1 and GFP::PRG-1 in the 4-cell embryos (left) or adult germline (right) of the indicated strains grown at their optimal growth temperature, 20°C. Arrow indicates the cytoplasmic P granules in the 4-cell embryo. (B) The brood size of worms of the indicated strains grown at 25°C. The standard error of the mean (SEM) are provided. *** indicates p< 0.001 using one-way ANOVA with Bonferroni multiple comparison test comparing indicated mutants to wild type animals. (C) A proposed model showing germline helicases couple their RNA binding and release to promote the assembly and disassembly of P granules. In the GLH-1 RNA binding mutant (middle), P granules fail to assemble and thus are reduced in size. On the contrary, in the GLH-1 mutant that is defective in RNA release (right), the P granule fails to disassembly and continues to assemble, leading to enlarged and less dynamic aggregates.

### Defects in perinuclear P granules correlate with fertility defects

As mentioned in the introduction, P granules are known to play a critical role in germ cell maintenance; various GLH mutants or GLH RNAi treated worms exhibit some fertility defects (Spike et al., 2008; Updike et al., 2014). We wonder whether the defects in fertility may correlate with the degree of P granule defects that are observed. We found a positive correlation between defects in forming perinuclear P granules and defects in fertility (Figure 4B). On the contrary, a previous study failed to identify a correlation between the degree of cytoplasmic P granule defects and infertility between several *meg* mutants (Wang et al., 2014). Furthermore, significant fertility defects were observed in GLH-1 FGGΔ mutant, which has abnormal perinuclear P granules but normal cytoplasmic P granules. Together, our observations suggest a critical role of perinuclear P granules in germ cell development.

## DISCUSSION

### Function of GLHs in regulating the dynamics of P granules

In this study, we have shown that germline helicases (GLHs) play multiple roles in controlling the dynamics and localization of P granules, the perinuclear liquid droplets found in the germlines of all animals. Our experiments support that GLHs, like its VASA homologue in fly, can act as an ATP-dependent RNA clamp, by which RNA binding and release is coupled to ATP binding and the release of ATP hydrolyzed products, respectively (Xiol et al., 2014). Importantly, we found that P granules fail to assemble in *glh* mutants that are defective in RNA binding, while P granules form enlarged aggregates in *glh* mutants defective in releasing RNAs. These observations suggest a simple model that the binding and release of RNAs from GLH-1 are coupled to the assembly and disassembly of P granules, respectively (Figure 4C). Our findings provide a simple explanation for the previous seemingly opposite effects of the RNA helicase VASA in germ granules in flies. For example, the VASA DQAD mutation also leads to enlarged germ granules in insect cells, while mutation at the conserved Threonine of motif V leads to loss of germ granules in *Drosophila* (Dehghani and Lasko, 2015; Xiol et al., 2014). Together, these observations suggest VASA-like helicases share a conserved role in regulating dynamics of germ granules in diverse animals. Importantly, DEAD box RNA helicases have also been reported to play a role in phase separation of other cellular bodies as well. For example, the mutation of DQAD in the yeast DHH1 helicase also leads to enlarged P bodies, while mutations affecting the RNA binding of DHH1 lead to loss of P bodies in yeast (Mugler et al., 2016). Together, our study provides an example of how RNA helicases control the formation and localization of phase-separated bodies and strengthen the emerging theme of DEAD box helicase as a key regulator of phase separation in cells (Hondele et al., 2019).

### GLH-1 drives the formation of P granules at perinuclear foci

Germ granules are frequently found at the perinuclear region in various animals. The mechanisms by which germ granules associate with the nucleus were not clear. In this study, we found that the FGG repeats of GLH-1 are critical for the formation of perinuclear P granules. In the GLH-1 FGGΔ mutant, GLH-1 no longer localizes to the nuclear periphery and the localization of piRNA factors to P granules is compromised. These observations are consistent with the model that the FGG repeats in GLH-1 interact with the hydrogels formed by FG-containing nucleoporins to anchor the helicase and P granules to the nuclear pores. Most VASA proteins in other animals do not have FGG repeats, but rather have Arginine-Glycine-Glycine RGG repeats, suggesting a distinct mechanism for the localization of VASA in germ granules. Intriguingly, the arginines of the VASA RGG repeats have been reported to be methylated, and the arginine methylation of RGG repeats of Aub Argonaute is essential for its localization to germ granules in the fruit fly (Kirino et al., 2010b, 2010a; Webster et al., 2015). Therefore, it is possible that various N terminal repeats in VASA helicases have evolved with different strategies to promote perinuclear localization of germ granules in animals.

### A potential role for P granule as a site for mRNA surveillance that promotes germ cell development

A previous study has demonstrated that perinuclear P granules are major sites of mRNA export, raising the intriguing hypothesis that P granules may facilitate mRNA surveillance (Sheth et al., 2010). Several observations made in our study support this model: First, we have shown that the inhibition of RNA Polymerase II transcription leads to the disassembly of perinuclear P granules in wild type animals. This observation suggests that the maintenance of perinuclear P granules requires newly made mRNAs and implies that GLHs associate with newly exported mRNAs to promote P granule formation. Second, our proteomic analyses showed that the interactions between GLH-1 and Argonautes are stabilized in the GLH-1 DQAD mutant, where RNA release is inhibited (Xiol et al., 2014), implying their interactions are mediated through RNAs and GLH-1 may facilitate the binding of Argonautes to their GLH bound mRNAs. Third, the degree of fertility defects in various VASA mutants correlates with the severity of defects in forming perinuclear P granules, but not the defects in forming cytoplasmic P granules, suggesting critical roles for perinuclear P granules in promoting germ cell viability. Taken together, our study provides strong implications that perinuclear P granules are poised to reinforce RNA regulation through Argonaute proteins. The extent to which P granule surveillance functions through small RNA-mediated gene regulation will be an important factor to investigate in the future.

## MATERIALS AND METHODS

### *C. elegans* strains

Animals were grown on standard nematode growth media (NGM) plates seeded with OP50 E. coli at 20°C or temperatures where indicated. The strains used in this study are listed in Table S1. Some strains were provided by the Caenorhabditis Genetics Center (CGC).

### RNAi

RNAi experiments were performed as described previously (Timmons and Fire, 1998). L1 hermaphrodites were fed with *E. coli* HT115 (DE3) strains expressing the appropriate double-stranded RNA (dsRNA) at 20°C for 72 hours prior to score for phenotypes. RNAi bacterial strains were obtained from the *C. elegans* RNAi Collection (Ahringer; Source BioScience). HT115 (DE3) strain expressing the empty vector L4440 was used as the control. HT115 (DE3) bacterial cultures were grown in Luria broth with 100 μg/ml ampicillin overnight at 37°C. Fresh cultures were seeded on NGM plates containing 100 μg/ml ampicillin and 1 mM IPTG and incubated at room temperature for 24 hours before use.

### CRISPR/Cas9 gene editing

sgRNAs were designed with the on-line tool sgRNA Scorer 2.0 (https://crispr.med.harvard.edu/), or manually picked by searching target sequences consisting of N_20_(NGG) which are most closed to DSB cutting sites desired. CRISPR experiments were conducted by co-CRISPR or Cas9 RNP strategies (Dokshin et al., 2018; Kim et al., 2014).

For co-CRISPR strategy, DNA mixtures were introduced into the germline of *C. elegans* young adults by microinjection. Final concentrations of plasmids in injection mixtures are as follows: pCCM935 unc-22 sgRNA 50 ng/μl, pDD162 Cas9+sgRNA 50 ng/μl, pRF4 rol-6dm 30 ng/μl, ssDNA donor oligo 50 ng/μl (or plasmid based donor 50 ng/μl). F1 *unc-22* twitchers or *rol-6* rollers were picked, followed by genotyping of desired mutations. When applicable, transgenic strains were outcrossed to remove *unc-22* mutations.

For RNP strategy, we followed the protocol described in (Dokshin et al., 2018). Final concentrations of injection components are as follows: Cas9 250 ng/μl, tracrRNA 100 ng/μl, crRNA 56 ng/μl were mixed and incubated at 37°C for 10 minutes, then ssDNA donor oligo 110 ng/μl, pRF4 rol-6dm 50 ng/μl were added and injection mixtures were introduced into the germline of *C. elegans* young adults by microinjection. F1 progenies from plates containing rollers were picked and genotyped to identify desired mutations.

### Plasmid construction

#### (1) Csa9/sgRNA constructs

We used the on-line tool sgRNA Scorer 2.0 (https://crispr.med.harvard.edu/) to design sgRNAs. The sgRNAs were cloned into pDD162 (Dickinson et al., 2013) by overlapping PCR using pDD162 as PCR template and the appropriate primers (see sgRNA sequences used in this study in Table S2), Overlapping PCR products were inserted into pDD162 linearized with SpeI/BsrBI digestion by seamless ligation cloning extract (SLiCE) (Zhang and Glotzer, 2015).

#### (2) Donor constructs

To generate flag::mCherry::glh-1 donor construct, 800bp upstream and 500bp downstream of glh-1, and flag::mCherry coding sequences were amplified by PCR using N2 genomic DNA or plasmids containing flag::mCherry as templates, and the primers WJC049 and LB18, LB20 and WJC137, WJC110 and WJC111, and WJC136 and LB14.

To generate glh-1FGdel donor construct, 900bp upstream and 500bp downstream of glh-1FGdel, and FGdel flanking sequences were amplified by PCR using N2 genomic DNA as templates, and the primers WJC049 and WJC050, WJC051 and WJC052 (see Table S2 for PCR primer sequences). To generate flag::prg-1 donor construct, 500bp upstream and downstream of prg-1 sequence were amplified by primers LB05 and LB27, LB20 and LB25, LB26 and LB10, respectively. PCR fragments were inserted into pUC19 linearized with HindIII / KpnI digestion by SLiCE. Silent mutations were introduced in gRNA targeting sites by site-directed mutagenesis in donor constructs using Phusion High-Fidelity DNA Polymerases (Thermo Scientific).

### Fluorescence microscopy

GFP- and mCherry-tagged fluorescent proteins were visualized in living nematodes or dissected embryos by mounting young adult animals on 2% agarose pads with M9 buffer (KH_2_PO_4_ 22 mM, Na_2_HPO_4_ 42 mM, NaCl 86 mM) with 10-50 mM levamisole, or mounting one-cell embryos on 2% agarose pads by dissecting gravid hermaphrodites into egg salt buffer (HEPES pH 7.4 5 mM, NaCl 118 mM, KCl 40 mM, MgCl_2_ 3.4 mM, CaCl_2_ 3.4 mM). Fluorescent images were captured using a Carl Zeiss LSM 800 confocal microscope with a 40X objective. Images were processed and quantified in ImageJ.

### Fluorescence Recovery After Photobleaching (FRAP)

Photobleaching and fluorescence recovery of GFP::PRG-1 and mCherry::GLH-1 in germlines or early embryos were conducted and recorded using a Carl Zeiss LSM 800 confocal microscope with a Plan-Apochromat 63x/1.4 Oil DIC M27 objective, controlled by ZEN software. ZEN built-in FRAP module was used to perform FRAP experiments. Bleaching was performed using 100% laser power in the 488 and 553 channels for GFP::PRG-1 and mCherry::GLH-1, respectively. The region slightly larger than a single granule was selected, and fluorescence intensity in the whole region was bleached to below 20% of initial intensity before fluorescence recovery was allowed. Multiple granules were selected in each imaging filed, and 3 of them were imaged without photobleaching to serve as normalization control. Images were acquired every 1 second during a recovery phase of 120 seconds after bleaching.

Images were processed and quantified in ImageJ. The total fluorescence intensity was measured for areas containing bleached granules (I), unbleached granules (I^norm^), or areas without granules (I^bkg^) at each time point. Fluorescence recovery rates were calculated as F_n_=(I_n_-I^bkg^_n_)(I^norm^_i_/I^norm^_n_)/(I_i_-I^bkg^_i_), where n stands for time points after photobleaching, and i stands for initial phase before photobleaching. Recovery rates for all granules at different time points were calculated and plotted by using GraphPad Prism 6.0.

### RNA Pol II inhibition

Transcriptional inhibitor a-amanitin (50 µg/ml in water; Sigma-Aldrich A2263) were injected into gonads of young adults. Injected worms were allowed to develop for 5hrs and then mounted on 2% agarose pads in M9 buffer with 2.5 mM levamisole, Fluorescent images were captured using a Carl Zeiss LSM 800 confocal microscope with a 40X objective. Images were processed and quantified in ImageJ.

### Immunoprecipitation

100,000 staged worms were frozen in liquid nitrogen and stored at −80 °C. Worm pellets were resuspended in equal volume of IP buffer (20 mM Tris-HCl pH 7.5, 150 mM NaCl, 2.5 mM MgCl2, 0.5% NP-40, 80 U ml-1 RNaseOUT, 1 mM dithiothreitol (DTT) and protease inhibitor cocktail without EDTA and homogenized using MP Fastprep 24 benchtop homogenizer set to 6M/s and run for 3x40s, keeping the tubes on ice for 5min between runs. Lysates were clarified by spinning down at 30,000 g for 15 min. Supernatants were precleared with 100 ul protein G beads and incubated with 10 ul anti-flag M2 antibody for 1h and bind to protein G beads for 1h at 4 °C. Beads were washed with wash buffer (20 mM Tris-HCl pH 7.5, 200 mM NaCl, 2.5 mM MgCl2, 0.5% NP-40) for six times and then resuspended in TBS buffer for mass spectrometry or western blotting.

### Mass Spectrometry

Samples on immunoprecipitated beads were denatured with 8M urea in 50 mM Tris-HCl, reduced with 5 mM Tris (2-carboxyethyl) phosphine hydrochloride (TCEP) and alkylated with 10 mM chloroacetamide (CAM). 50 mM Tris-HCl, pH 8.5 was added to dilute urea to 2 M. 0.5 μg trypsin/LysC mix (Promega) was added overnight at 37°C. The supernatant was collected from all samples and cleaned up with C-18 spin columns (Thermo Fisher).

Thermo-Fisher Scientific Q Exactive Plus Mass Spectrometer coupled with Thermo-Fisher Scientific Easy-nLC1200 were used for analysis. Digested peptides were loaded onto an Acclaim PepMap C18 trapping column (3 μm particle size, 100 Å pore size, 2 cm length, 75 μm outer diameter) and eluted on a PepMapTM RSLC C18 column (2 μm particle size, 100 Å pore size, 15 cm length, 75 μm outer diameter) with a linear gradient from 3 to 35% acetonitrile (in water with 0.1% FA) developed over 120 min at room temperature at a flow rate of 650 nL/min, and effluent was electro-sprayed into the mass spectrometer. A blank was run prior to the sample run to make sure there was no significant signal from solvents or the column.

Raw files generated from the run were analyzed using Thermo Proteome Discoverer (PD) 2.2. SEQUEST HT was utilized to perform database searches with a few modifications: trypsin digestion, 2 maximum missed cleavages, precursor mass tolerance of 10 ppm, fragment mass tolerance of 0.8 Da, a fixed modification of +57.021 Da on cysteine, and a variable modification of +15.995 Da on methionine. The spectral false discovery rate (FDR) was set to ≤ 1%. The FASTA database used was a C.elegans proteome downloaded from Uniprot.

### West Blotting

Lysates were prepared from ∼100 synchronized young adult worms by boiling worms in 1x SDS sample buffer for 10 minutes. Proteins were separated by standard SDS-PAGE, and transferred to PVDF membranes (Millipore) using a Trans-Blot Turbo Transfer System (Bio-Rad). Membranes were blocked in TBST (20 mM Tris-HCl pH 7.4, 150 mM NaCl, 0.05% Tween-20) containing 5% skim milk for 2 hours at room temperature, then probed with primary antibodies against PRG-1 (1:1000, gift from C.C. Mello), GLH-1 (1:1000, gift from K.L. Bennett) and FLAG (1:1000, Sigma-Aldrich F1804) overnight at 4°C, separately. After 5 times washes with TBST buffer, the membranes were incubated with secondary antibodies against the species of primary antibodies at RT for 1hr and then washed 5 times with TBST, ECL substrates were used for detection of proteins by FUJIFILM LAS 4000 luminescent image analyzer.

### Ex vivo assay

Gravid young adult hermaphrodites were dissected in ex vivo buffer (10 mM Tris pH 7.5, 300 mM NaCl, 5 mM EDTA pH 7.5), and gonads were cut by a needle to let granules released into the buffer. When needed, 100 U/ml RNaseA/T1 mix (Thermo Scientific) was added into ex vivo buffer. Dissected gonads were then mounted on 2% agarose pads made with ex vivo buffer. Images were captured using a Carl Zeiss Axio Scope.A1 compound fluorescent microscope with an EC Plan-Neofluar 40x/0.75 M27 objective and a Retiga R6 CCD (Teledyne QImaging). Images were processed and quantified in ImageJ.

### Brood size analysis

Single hermaphrodite L4s (P0) were picked onto individual freshly-seeded NGM plates and allowed to grow for 24 hours at 20°C or 25°C. P0 adults were transferred to new NGM plates every 24 hours until they no longer laid eggs. All F1 progenies on each plate were counted. The brood size of each P0 animal is the total sum of F1s for all plates where the P0 animal laid eggs.

## ACKNOWLEDGEMENT

We thank Dr. Edwin Ferguson for the critical comments on the manuscript; members of the Lee lab for helpful discussions. Some strains used in this study were provided by the CGC, which is funded by NIH Office of Research Infrastructure Programs (P40 OD010440). This work is supported in part by NIH predoctoral training grant T32 GM07197 to J.B.; the National Natural Science Foundation of China (grants 31771500 and 31922019) and the program for HUST Academic Frontier Youth Team (grant 2018QYTD11) to D.Z; NIH grants R00-108866 and R01-GM132457 to H.-C.L.

---

**Figure S1.**
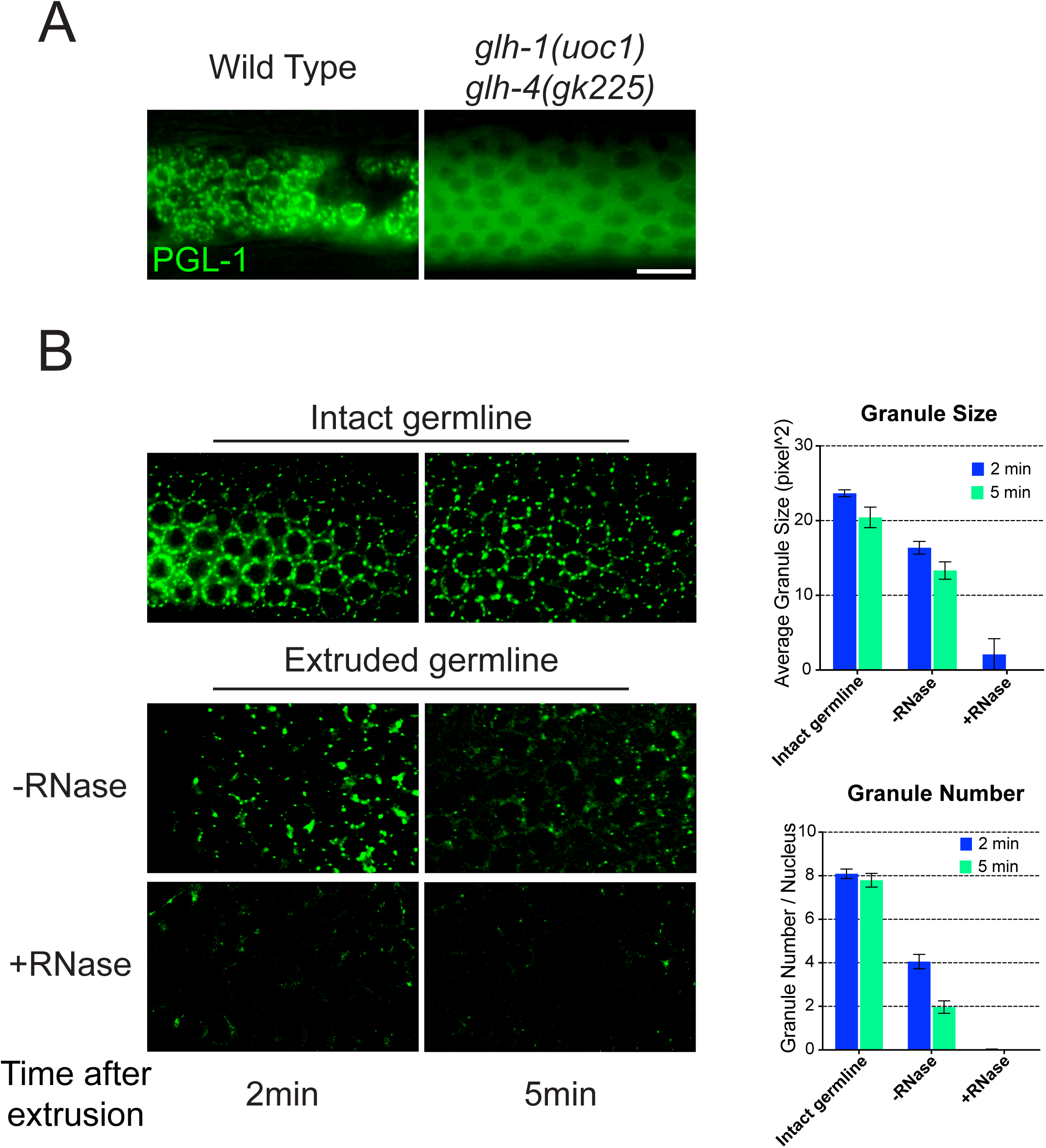
Germline helicases promote the formation of P granules. (A) GLH-1 and GLH-4 act redundantly to promote the formation of P granules. Fluorescent micrographs showing the localization of P granule component GFP::PGL-1 in the indicated strains. *uoc1* and *gk225* are null alleles of indicated genes. (B) RNAse treatment promotes the *ex vivo* disassembly of P granules. Fluorescent micrographs showing P granules at the indicated time after germlines were extruded in the RNase containing buffer or in the control buffer.

**Figure S2.**
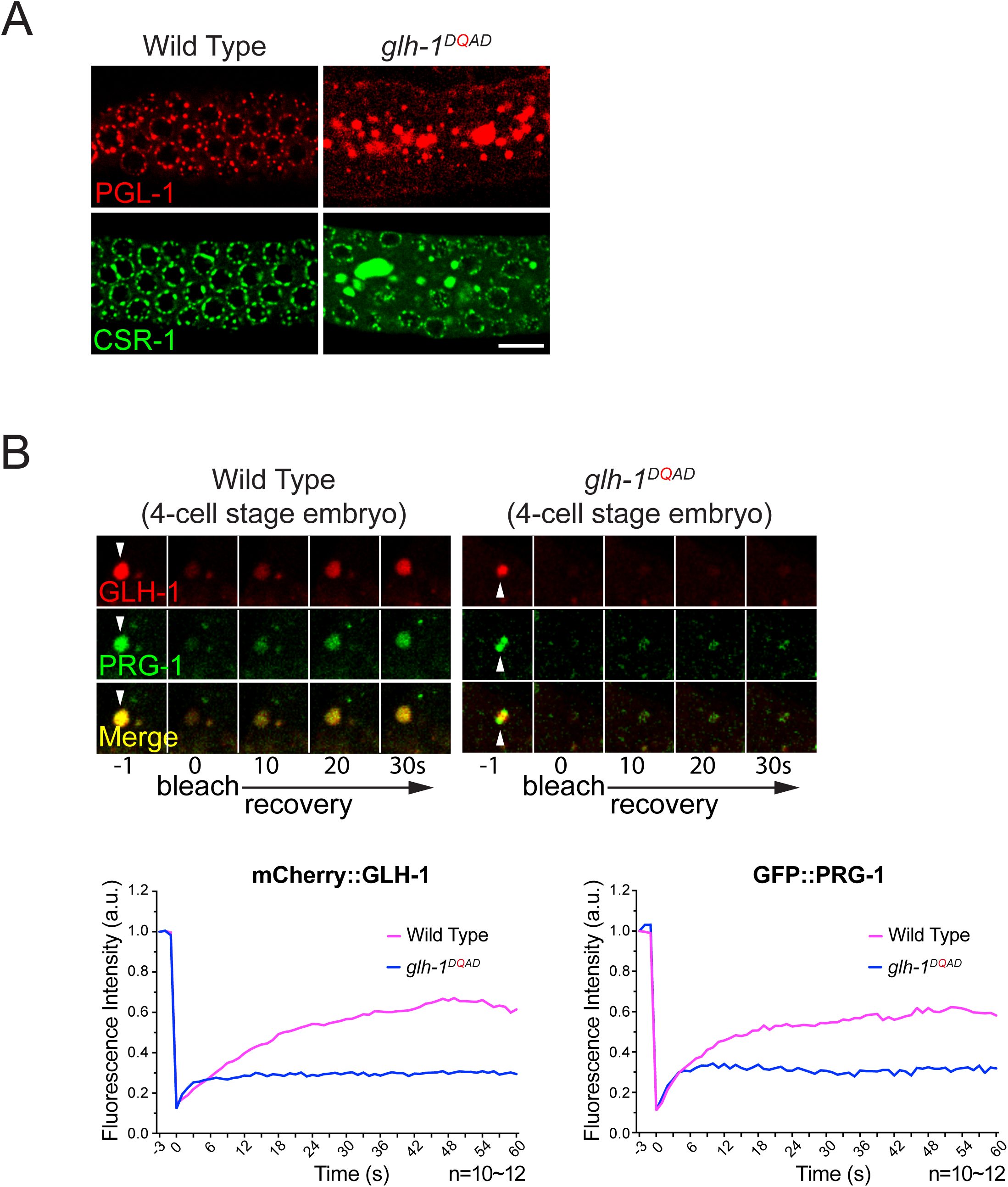
P granules form enlarged and less dynamic aggregates in *glh-1* DQAD mutant. (A) Fluorescent micrographs showing that components of P granules, such as mCherry::PGL-1, and GFP::CSR-1 are abnormally aggregated in the *glh-1* ^DQAD^ mutant. (B) The FRAP analyses indicate that the dynamics of P granule components, including mCherry::GLH-1 and GFP::PRG-1, are both reduced in *glh-1*^DQAD^ in the 4-cell embryos. The arrowheads indicate the P granules that are photobleached.

**Figure S3.**
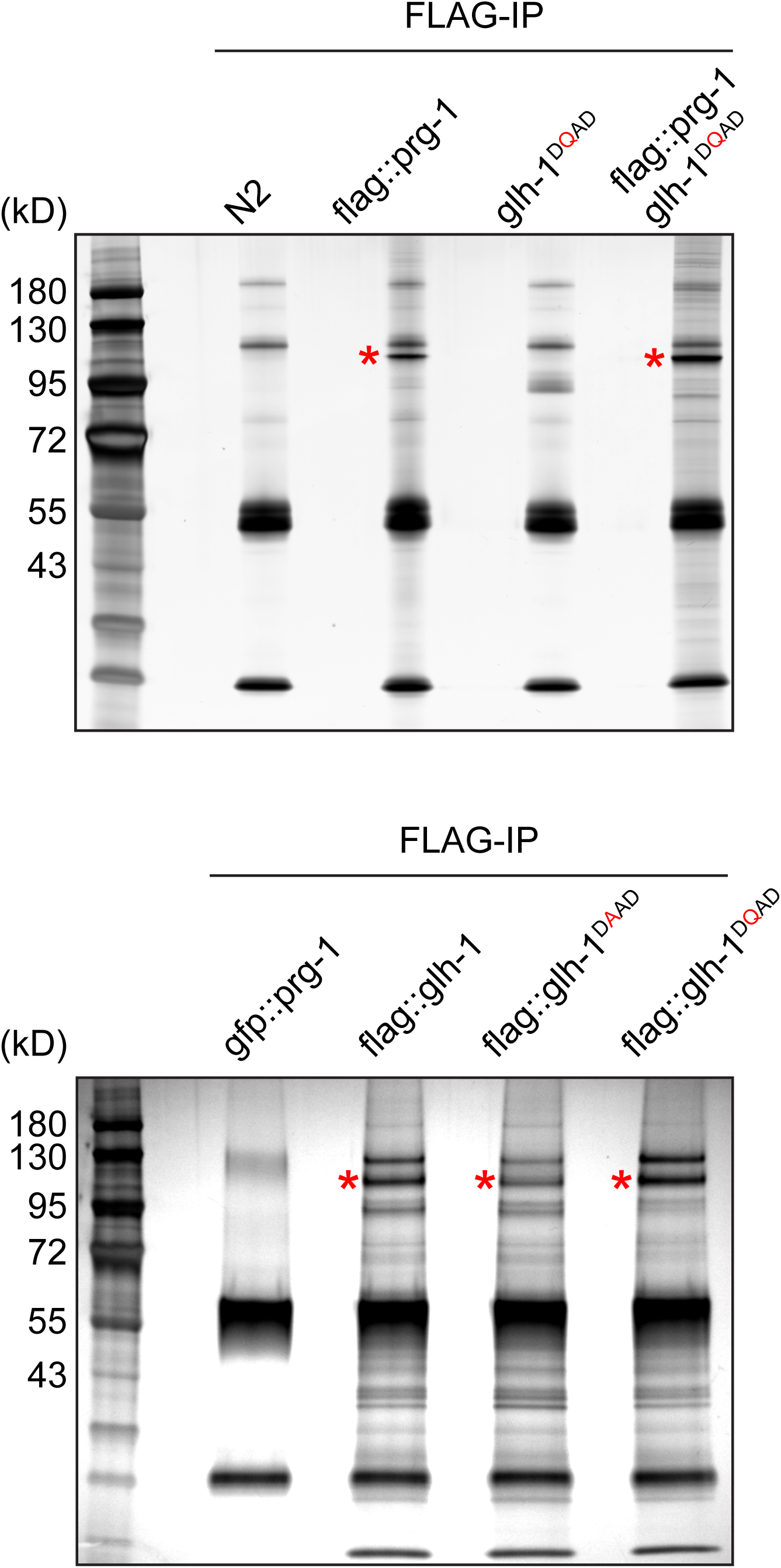
Proteomic analyses of GLH-1 and PRG-1 complex in *glh* mutants. Silver staining of PRG-1 complex (top) or GLH-1 complex (bottom) isolated from the adult worms of the indicated strains. Asterisks indicate the expected positions of FLAG::PRG-1 (top) and FLAG::GLH-1 (bottom), respectively.

**Figure S4.**
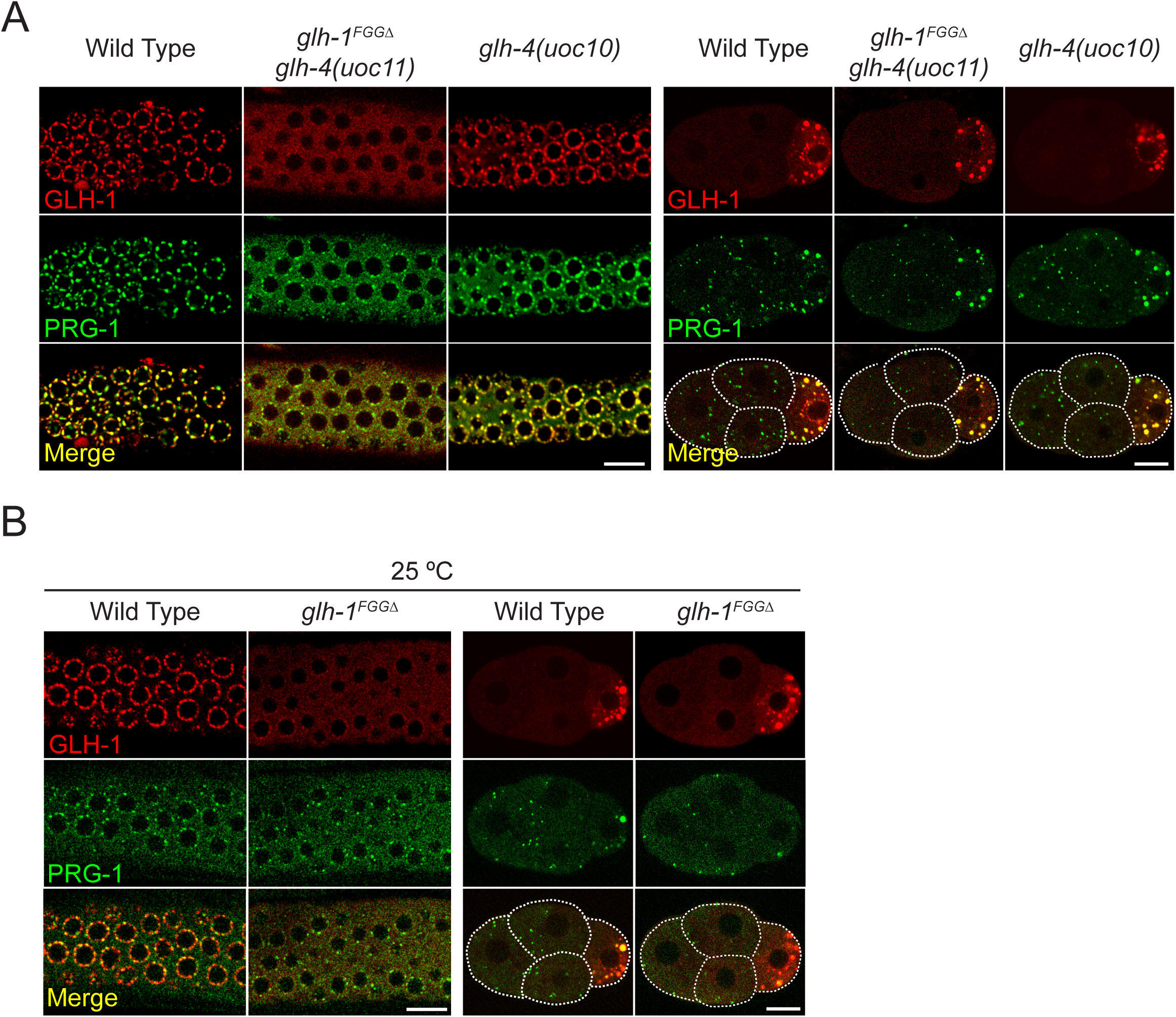
The N terminal FGG repeats in GLH-1 promote nuclear anchoring of P granules. (A) The formation of perinuclear P granules in the adult germline, but not cytoplasmic P granules in the embryos, is compromised in the *glh-1*^FGGΔ^ *glh4* double mutant. Images showing the localization of mCherry::GLH-1 and GFP::PRG-1 in the adult germline (left) or the four-cell embryo (right) of the indicated strains grown at their optimal growth temperature, 20°C. (B) The formation of perinuclear P granules in the adult germline, but not cytoplasmic P granules in the embryos, is compromised in the *glh-1*^FGGΔ^ mutant. This image shows the localization of mCherry::GLH-1 and GFP::PRG-1 in the adult germline (left) or the four-cell embryo (right) of the indicated strains grown at 25°C.

**Table S1.** C. elegans strains used in this study

**Table S2.** Oligonucleotide sequences used in this study

**Table S1.** *C. elegans* strains used in this study.

| Strain | Genotype | Method |
| --- | --- | --- |
| HCL117 | *gfp::prg-1 glh-4(uoc10) I* | CRISPR/Cas9 |
| HCL118 | *glh-1(uoc1) I* | CRISPR/Cas9 |
| HCL119 | *glh-1<sup>DAAD</sup>(uoc2) I* | CRISPR/Cas9 |
| HCL120 | *glh-1<sup>DQAD</sup>(uoc3) I* | CRISPR/Cas9 |
| HCL121 | *glh-1<sup>FGGΔ</sup>(uoc4) I* | CRISPR/Cas9 |
| HCL122 | *glh-1(uoc5) glh-4(gk225) I/hT2 [bli-4(e937) let-?(q782) qIs48] (I;III)* | CRISPR/Cas9 |
| HCL123 | *gfp::prg-1 glh-1(uoc1) I* | Cross |
| HCL124 | *gfp::prg-1 glh-1(uoc1) glh-4(uoc6) I* | CRISPR/Cas9 |
| HCL125 | *gfp::prg-1 flag::mCherry::glh-1 I* | CRISPR/Cas9 |
| HCL126 | *gfp::prg-1 flag::mCherry::glh-1<sup>DAAD</sup> I* | CRISPR/Cas9 |
| HCL127 | *gfp::prg-1 flag::mCherry::glh-1<sup>DQAD</sup> I* | CRISPR/Cas9 |
| HCL128 | *gfp::prg-1 flag::mCherry::glh-1<sup>FGGΔ</sup> I* | CRISPR/Cas9 |
| HCL129 | *gfp::csr-1 IV; glh-1(uoc1) I* | Cross |
| HCL130 | *gfp::csr-1 IV; glh-1<sup>DQAD</sup>(uoc3) I* | Cross |
| HCL131 | *gfp::csr-1 IV; glh-1(uoc5) glh-4(gk225)* | Cross |
| HCL132 | *gfp::pgl-1 IV; glh-1<sup>DQAD</sup>(uoc3) I* | Cross |
| HCL133 | *gfp::pgl-1 IV; glh-1(uoc5) glh-4(gk225) I* | Cross |
| HCL143 | *gfp::prg-1 flag::mCherry::glh-1(T663A) I* | CRISPR/Cas9 |
| HCL150 | *gfp::prg-1 flag::mCherry::glh-1<sup>FGGΔ</sup> I glh-4(uoc11)* | CRISPR/Cas9 |

**Table S2.**
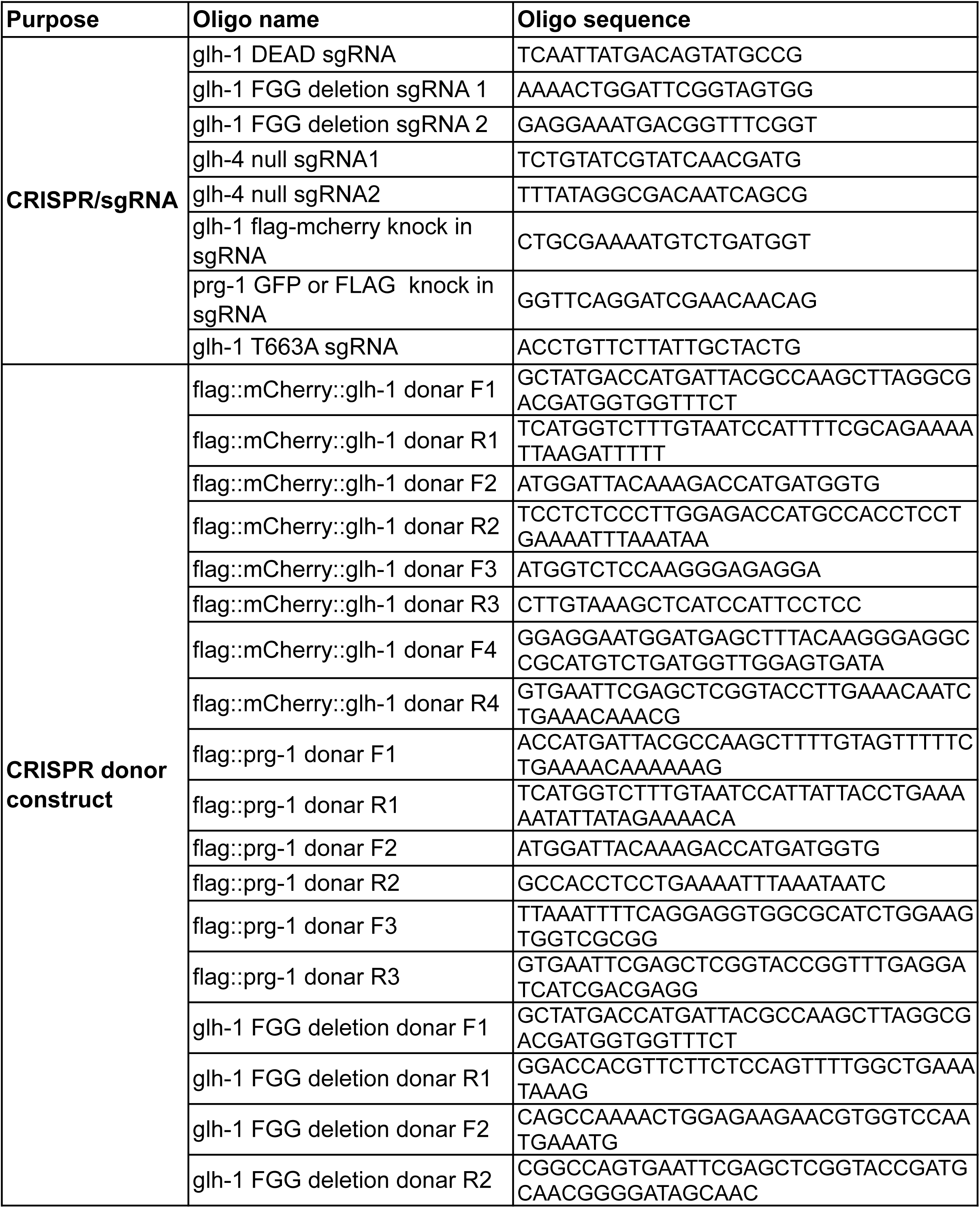
Oligonucleotide sequences strains used in this study.

